# Glycerol Production and Diapause Are Most Likely Not Linked in *Manduca sexta*

**DOI:** 10.1101/2025.04.29.649890

**Authors:** Robert Ades

## Abstract

Several studies have demonstrated that diapausing insects upregulate glycerol production in response to cold temperatures. However, relatively few studies have investigated whether diapause can independently trigger increased glycerol production. In other words, little research has been dedicated to analyzing whether diapause and glycerol upregulation are linked. If a linkage between these two elements in insects exists, then photoinduced diapausing insects should upregulate glycerol production, even without exposure to cold temperatures. The current study examines whether this linkage exists in the tobacco hornworm pupae (*Manduca sexta*). Tobacco hornworms will enter a pupal diapause if presented with consistently short photoperiods, most notably 12 hours of daylight followed 12 hours of darkness. Regarding temperature, photoinduction of diapause in the tobacco hornworm is optimal at 26 °C. Using a spectrophotometric assay, glycerol levels in hemolymph were assessed in non-diapausing pupae shortly after pupation and in diapausing pupae after 16 days of pupation. Interestingly, non-diapausing pupae contained significantly higher glycerol levels in hemolymph compared to diapausing pupae. Furthermore, non-diapausing pupae produced approximately .099 M glycerol and diapausing pupae produced .085 M glycerol. This study provides evidence that glycerol upregulation and diapause are not linked in *Manduca sexta*, but are rather two separate events with two distinct causes. Nevertheless, the results are valuable since non-diapause specimens produced more glycerol than diapausing specimens, and moderately high concentrations of glycerol were found in both diapausing pupae and non-diapausing pupae.

---

Glycerol is a biomolecule that is synthesized in many organisms, including insects, animals, humans, and plants (Araújo et al., 2019; Fraser et al., 2017; Gerber et al., 1988; Ishiguro et al., 2007; Nelson et al., 2011). However, different organisms produce glycerol for a wide array of reasons. For example, in humans and animals, glycerol is a substrate for gluconeogenesis, a component of many metabolic processes, and a precursor for synthesizing triacylglycerols and phospholipids (Leithner et al., 2018; Rotondo et al., 2017). These functions indicate that humans use glycerol to produce energy sources and structural support for cell membranes. Similarly, glycerol is involved in metabolic processes for plants, but several studies have also demonstrated that plants use glycerol-3-phosphate, a phosphorylated form of glycerol, for defense against certain pathogens (Chanda et al., 2017; Yang et al., 2013). Interestingly, in addition to the several possible functions of glycerol in insects, some insects uniquely use glycerol as a cryoprotectant to survive cold temperatures. Specifically, the goldenrod-gall fly, false darkling beetle, rice-stem borer, and a few other insects upregulate glycerol production for overwintering because of glycerol’s cryoprotective properties (Dubach et al., 1959; Ishiguro et al., 2007; Rickards et al., 1987). Primarily, glycerol competes for hydrogen bonds on water molecules, consequently disrupting intermolecular forces between water molecules and suppressing the formation of ice (Grasso et al., 2018). Furthermore, glycerol stabilizes macromolecules, cells, and tissues that are below freezing temperatures (Grasso et al., 2018). Although several studies signify that many diapausing insects upregulate glycerol production to survive cold temperatures, relatively few studies have investigated whether diapause can exclusively trigger increased glycerol production. Moreover, limited research is available on whether *Lepidoptera* insects produce glycerol for cold-hardiness, and no study has been conducted to investigate if the *Lepidoptera* family *Sphinigidae* uses glycerol for cryoprotection. The current study will examine whether *Manduca sexta*, otherwise known as the tobacco hawkmoth or tobacco hornworm, will increase glycerol production during diapause. Diapause is defined as a period when an organism enters a metabolic and developmental arrest, usually because of unfavorable environmental conditions (Delinger et al., 2009). In *Manduca sexta*, diapause may only occur during the pupal stage, and will normally transpire in response to short photoperiods. Many insects, including the tobacco hornworm, recognize that short photoperiods imply that the onset of winter is temporally close and approaching (Bell et al., 1975).

In a landmark study on this topic, Rickards et al. (1987) investigated glycerol production in the goldenrod gall fly larva. First, Rickards et al. (1987) obtained goldenrod gall fly larvae from goldenrod fields in Ottawa, Canada. Once temperatures started to drop significantly, the researchers placed the larvae outdoors to expose the specimens to sustained winter temperatures. Rickards et al. (1987) periodically collected samples of these larvae from November to May, in attempt to reveal how glycerol levels change during the winter and spring. Remarkably, the results displayed a significant increase in glycerol levels during the winter months, directly in line with the decrease in outdoor daily temperature. Furthermore, after winter transitioned to spring, glycerol levels rapidly decreased in a manner closely in line with the increase in outdoor daily temperature. These results denote that glycerol acts as an antifreeze in goldenrod gall fly larvae, and the larvae amplify production of this antifreeze during the cold months to assist with overwintering (Rickards et al., 1987).

In another notable study on this topic, Lee et al. (1987) examined whether inducing pupal diapause in flesh flies (*Sarcophaga crassipalpis*) will lead to a larger production of cryoprotectants. In particular, this study photoinduced diapause in one group of flesh flies, and reared this group of flesh flies at room temperature. Photoinduction of diapause is defined as the onset of diapause in response to consistently short photoperiods, typically around 12 hours of daylight and 12 hours of darkness. Instead of placing these flies in cold temperatures, the researchers placed these specimens in room temperature to unveil whether diapause can independently trigger the upregulation of cryoprotectants without exposure to cold temperature. Essentially, the cryoprotectants of these flesh flies that were in diapause were compared to a group of flesh flies that were not in diapause and raised at the same temperature. Strikingly, Lee et al. (1987) observed that diapausing pupae had significantly greater cold tolerance than non-diapausing pupae. Furthermore, although non-diapausing pupae and diapausing pupae had similar glycerol levels in the beginning of the first couple of weeks of the pupal stage, the diapausing pupae experienced a steady increase in glycerol levels for nearly a month. Moreover, at the termination of diapause, the glycerol concentration in the diapausing pupae rapidly declined (Lee et al., 1987). These results provide evidence that diapause and cryoprotectant production are directly linked in some insects.

Although Lee et al. (1987) provided an example of a direct relationship between diapause and glycerol production, most previous studies conducted on this topic do not support this direct linkage. Instead, the vast majority of insects studied demonstrate that exposure to cold temperatures is required for enhanced glycerol production, and therefore diapause cannot independently increase glycerol production. This determination was found in the European corn borer (*Ostrinia nubialis*; Hanec & Beck, 1960), the viceroy butterfly (*Limentis archippus*; Frankos & Platt, 1976), and two springtail insects (*Orchesella cincta*; *Tomocercus minor*; Woude & Verhoef, 1988). Additionally, in some insects, low temperature exposure to non-diapausing insects can increase glycerol production (Wood & Nordin, 1976).

The current study tested for the potential linkage between diapause and glycerol production in tobacco hornworm (*Manduca sexta*). Additionally, this study analyzed glycerol levels during the pupal stage of tobacco hornworms because diapause may only occur during the pupal stage in this insect. Two groups were involved in this study: one group of tobacco hornworms that were photoinduced into diapause, and a non-diapause group of tobacco hornworms that were reared at normal light and temperature conditions. Hemolymph samples were taken from the non-diapause group of tobacco hornworms shortly after pupation. For the diapause group, hemolymph samples were obtained sixteen days after pupation. In order to measure glycerol in the blood samples, a Beer’s Law plot was initially created using known concentrations of glycerol and their measured absorbances. Subsequently, a spectrophotometric glycerol assay invented by Sturgeon et al. (1979) was used to measure glycerol concentrations in the collected hemolymph samples. Importantly, this same assay was used for the Beer’s Law plot samples. The null hypothesis for this experiment claims that no differences in glycerol production will be evident between the diapausing and non-diapausing pupae. However, some research supports an alternative hypothesis, which claims that the diapausing specimens will produce higher glycerol concentrations than the non-diapausing specimens. This study is valuable for a few main reasons. The first reason is that *Manduca sexta* is an agricultural pest, and an extensive understanding of diapause in *Manduca sexta* may assist with monitoring the populations of this agricultural pest. As noted in many studies, the tobacco hornworm is a pest of several crops, including tomato, pepper, tobacco, potato, and eggplant (Byron & Gillet-Kaufman, 2018). Secondly, many studies have measured glycerol hemolymph in a variety of insects, but no study has been conducted on glycerol hemolymph levels in the tobacco hornworm. This study provides insight into glycerol hemolymph levels of both diapausing and non-diapausing tobacco hornworms. Lastly, few studies have uncovered a link between diapause and glycerol production because most insects exhibit no link between dormancy and cryoprotectant production. The current study investigates whether this relationship between diapause and glycerol production exists in *Manduca sexta*.

## Method

### Insect Care and Subjects

Originally, the experiment reared 15 tobacco hornworms in each group, for a total of 30 tobacco hornworms. Unfortunately, 3 specimens from each group either died during rearing or failed to pupate, leaving 12 tobacco hornworms in each group (*n* = 12) and a total of 24 specimens (*N* = 24). All tobacco hornworms were initially reared in small individual containers, and moved to larger individual containers after reaching the third instar. All specimens were ensured to have adequate food every other day. Additionally, if any fungus was observed on a food sample, then the spoiled piece of food would be replaced with a fresh piece. The food diet from Bell and Joachim’s (1976) study was used for rearing all the tobacco hornworms. Frass, or fecal matter, was also cleaned every other day from the containers.

### Materials

This experiment used a between-subjects design with one control group (*n* = 12) and one experimental group (*n* = 12). The control group was the non-diapause group that was reared under normal light and temperature conditions. Conversely, the diapause group was the experimental group, which was reared in an incubator at alternate light and temperature conditions to induce diapause. Therefore, the independent variable was whether the specimens were induced into diapause. Centrifuge tubes and utility blades were used to collect the original supernatant samples, and glycerol concentrations in µg/mL were measured. Accordingly, glycerol concentration in µg/mL was the dependent variable for the experiment. Regarding the glycerol assay, many materials were needed, including a .5-10 µL pipette, stainless steel pot, Bunsen burner, 100 mL volumetric flasks, 10 mL graduated cylinders, 10 mL pipettes, 16 × 100 millimeter (mm) test tubes, and a spectrophotometer with cuvettes. The stainless steel pot, Bunsen burner, and volumetric flasks were used to heat the solutions in a 100 °C water bath, while the spectrophotometer and cuvettes were needed to measure the absorbances of each solution at 570 nanometers (nm). In addition, several reagents were required, including glycerol (C_3_H_8_O_3_), sodium periodate (NaIO_4_), sodium sulfite (Na_2_SO_3_), chromotropic acid (C_10_H_8_O_8_S_2_), sulfuric acid (H_2_SO_4_), and distilled water (H_2_O).

### Procedure

#### Insect Rearing

All specimens were first reared in individual plastic containers until the fifth instar. The non-diapause group was exposed to light-dark cycles of 16 hours light followed by 8 hours dark (16L:8D) at room temperature, which was approximately 23°C. On the other hand, the diapause group was placed in an incubator that was programmed to provide a light cycle of 12 hours light followed by 12 hours dark (12D:12L). The incubator was set to 26°C, as this temperature falls in the optimal range for photoinduction of diapause in *Manduca sexta* (Bell et al., 1975). This photoinduction of diapause method was taken from Bell et al.’s (1975) study, which established that *Manduca sexta* will enter a pupal diapause if the larvae are exposed to 12L:12D cycles at 26°C. Once the specimens clearly underwent the wandering stage of development, they were placed into cylindrical holes of wooden blocks to promote pupation.

### Collection of Hemolymph Samples

Immediately after the non-diapause specimens pupated, hemolymph samples were obtained. Concerning the diapausing pupae, hemolymph samples were acquired 16 days after pupation. In order to collect a hemolymph sample, an incision was made in the wing of each pupa, directly under the antenna (Fig. 1). The hemolymph would typically spill out of the incision, and the supernatant was collected in centrifuge tubes. Immediately after collecting a specimen’s supernatant, the specimen would be put in the freezer to minimize suffering.

**Figure 1.**
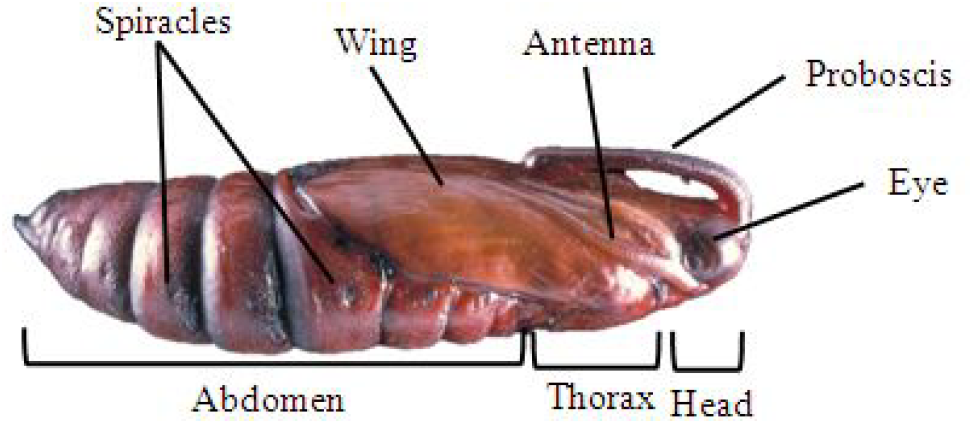
*Manduca sexta* Pupa Anatomy (Berkley, 2019)

### Glycerol Assay

In this glycerol assay, glycerol is first oxidized by sodium periodate to form formaldehyde. Next, the formaldehyde reacts with chromotropic acid to produce a colored product. For illustration purposes, these are the following reactions:

1. Glycerol + sodium periodate → formaldehyde + formic acid
2. Chromotropic acid + formaldehyde → colored product

For the second reaction to occur, excess acid must be present, which will be provided in the form of sulfuric acid.

Specifically, these were the exact instructions for the glycerol assay from Sturgeon et al. (1979): First, pipette a 2.0 mL sample containing anywhere between 0–1000 µg of glycerol/mL into a 16 × 1000 mm test tube. Next, add 1 mL of 1% NaIO_4_, mix the sample in the test tube, and leave the solution standing at room temperature for 15 minutes. Subsequently, add 1 mL of 5% Na_2_SO_3_ to the mixture, and allow this solution to stand for 5 minutes at room temperature. Using a pipette, transfer 1.0 mL of this mixture into a 100 mL volumetric flask, and add .50 mL of 10% chromotropic acid to this mixture. 5.0 mL of concentrated H_2_SO_4_ will then be added to the mixture in the flask, and should be added slowly to prevent any explosion from occurring. Next, heat this solution in a 100 °C water bath for 30 minutes. After heating, add 10 mL of distilled water and allow the solution to cool to room temperature. Lastly, using a cuvette and spectrophotometer, measure the absorbance of the solution at 570 nm (Sturgeon et al., 1979).

A Beer’s Law plot was created using this glycerol assay with known concentrations of glycerol. The exact concentrations of glycerol used for this plot, in units of µg/mL, were 25, 50, 75, 100, 150, 200, 250, 300, 350, 400, 450, and 500. A linear equation was obtained for the Beer’s Law plot, which was used to calculate the glycerol concentrations in the hemolymph samples.

One important note should be made regarding this glycerol assay, as pertaining to the hemolymph samples. The first step of this glycerol assay is to pipette a 2.0 mL sample containing anywhere between 0–1000 µg/mL of glycerol in a 16 × 100 mm test tube. However, each specimen only provided .15–.25 milliliters of hemolymph. To account for this issue, .1 mL of hemolymph was used from each sample and the volume was raised to 2.0 mL with distilled water. After using the Beer’s Law plot equation to obtain the measured concentrations, each measured concentration was multiplied by a factor of 20 to correct for the initial dilution. In other words, the hemolymph solutions were originally diluted by a factor of 20 in the assay, and the measured glycerol concentrations were accordingly multiplied by a factor of 20 to obtain the corrected glycerol concentrations.

## Results

The original Beer’s Law plot displayed a relatively exponential design (Fig. 2), since the function was not linear for glycerol concentrations below 200 µg/mL (*r* = .934, *r*^*2*^ = .872). Instead of this plot, a new Beer’s Law plot was created with only the glycerol concentrations of 200 µg/mL and above (Fig. 3), since these concentrations are included in the linear range for the assay (*r* = .983, *r*^*2*^ = .967). This new Beer’s Law Plot produced an equation of y = .0036x – .5786, and this equation was used to calculate the glycerol concentration in each specimen’s hemolymph. Both the original Beer’s Law plot (Fig. 2) and linear range Beer’s Law plot (Fig. 3) are provided for transparency. From the hemolymph samples, all measured glycerol absorbances were within the linear range of the Beer’s Law plot, which ensures accuracy. An independent-measures two-tailed t-test was used to assess the potential differences in glycerol concentrations between non-diapausing pupae and diapausing pupae. For this t-test, the alpha level used was .05. The non-diapausing pupae (*µg/mL* = 9079.7, *SD* = 681.4) reported significantly higher glycerol concentrations (Fig. 4) than the diapausing pupae (*µg/mL* = 7814.4, *SD* = 1259.4), [*t*(22) = 3.06, *p* = .0057]. In terms of molarity, the diapausing specimens produced an average of .085 M and the non-diapausing specimens produced an average of .099 M.

**Figure 2.**
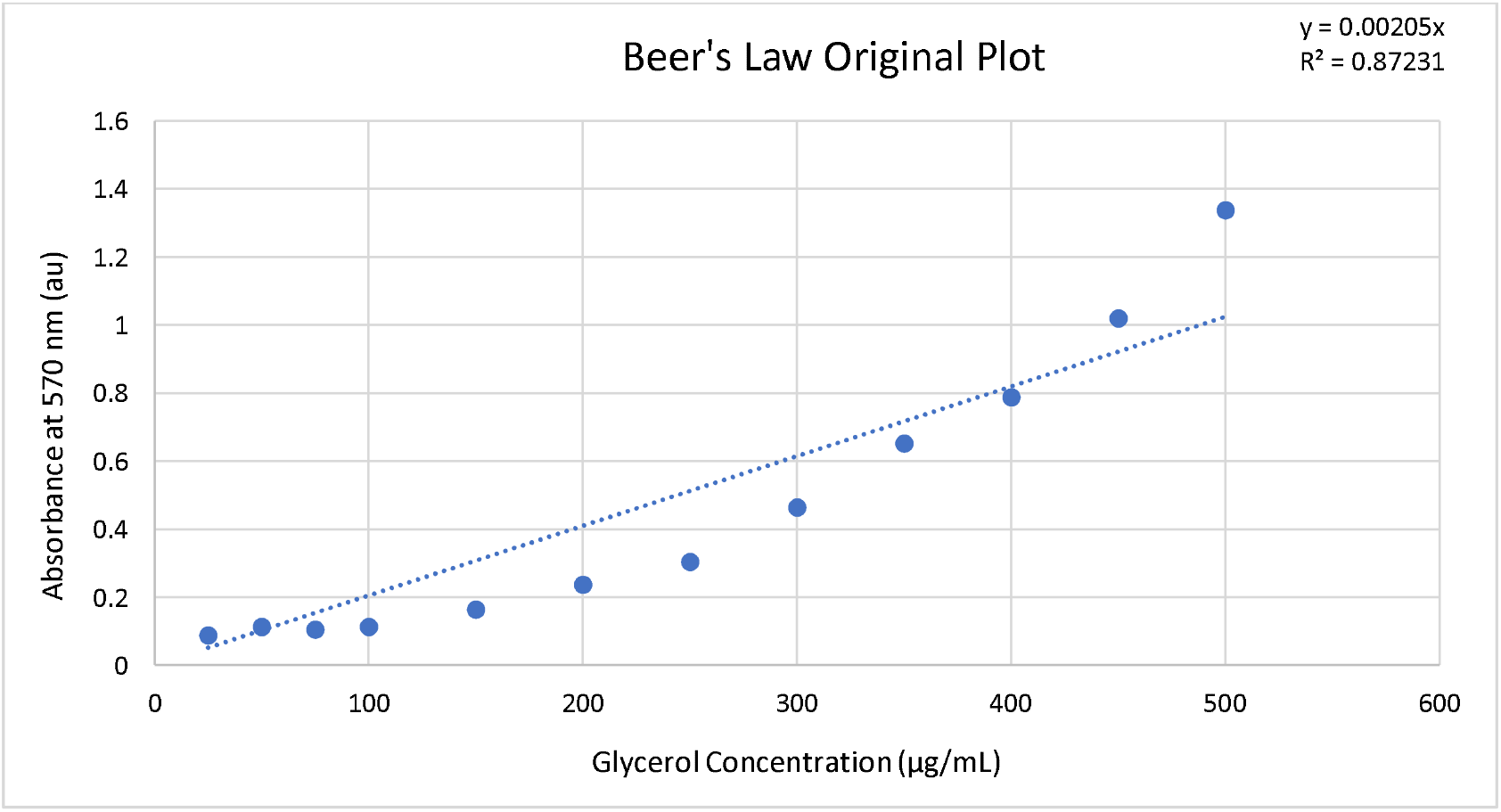
The original Beer’s Law plot performed with exact concentrations of glycerol, in µg/mL, of 25, 50, 75, 100, 150, 200, 250, 300, 350, 400, 450, and 500.

**Figure 3.**
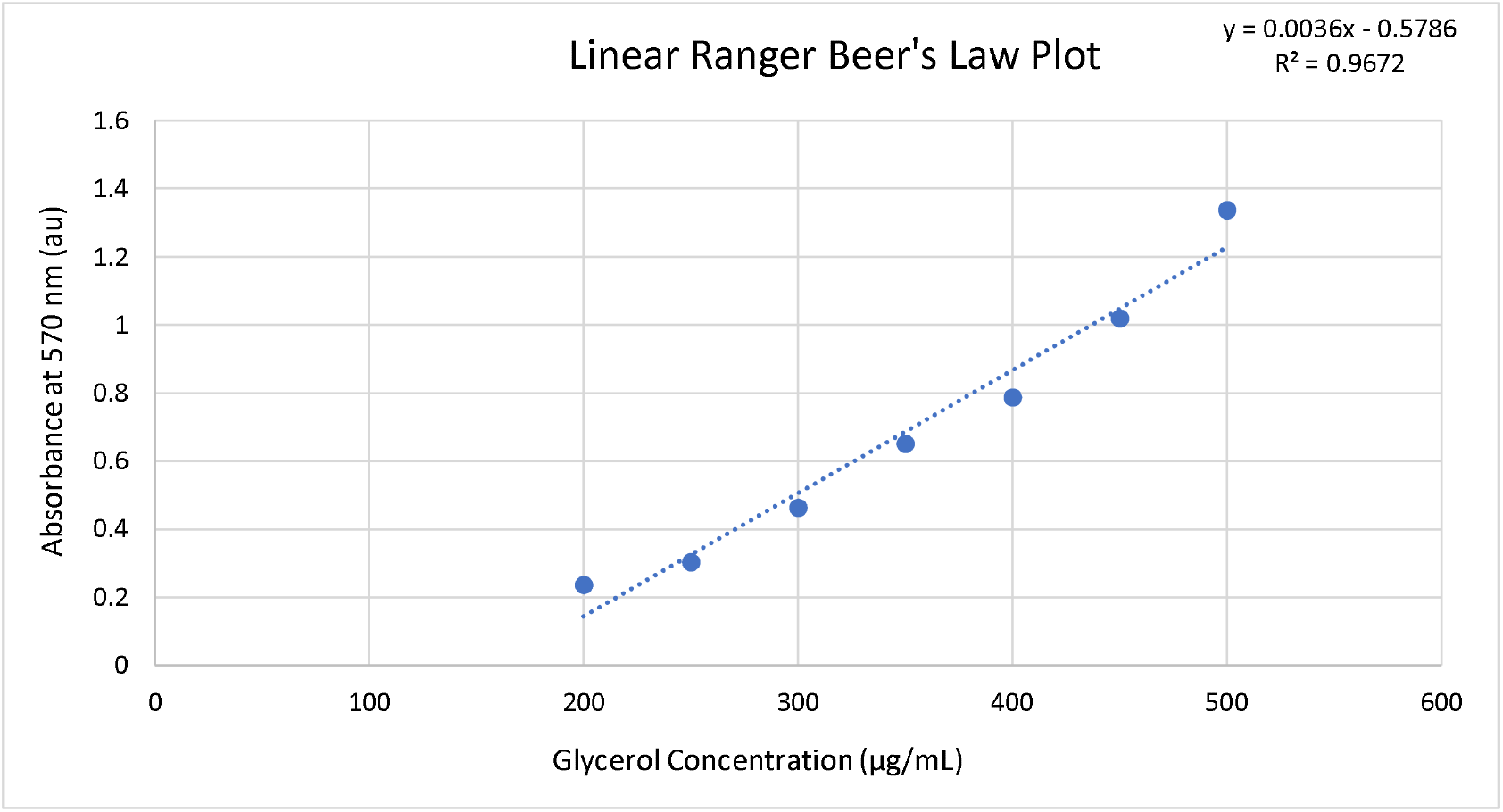
Illustrated here is the linear range of the Beer’s Law plot, consisting only of glycerol concentrations from 200–500 µg/mL (200, 250, 300, 350, 400, 450, and 500 µg/mL). Glycerol concentrations below 200 and above 500 were excluded from the original Beer’s Law plot to obtain the linear range depicted in this plot.

**Figure 4.**
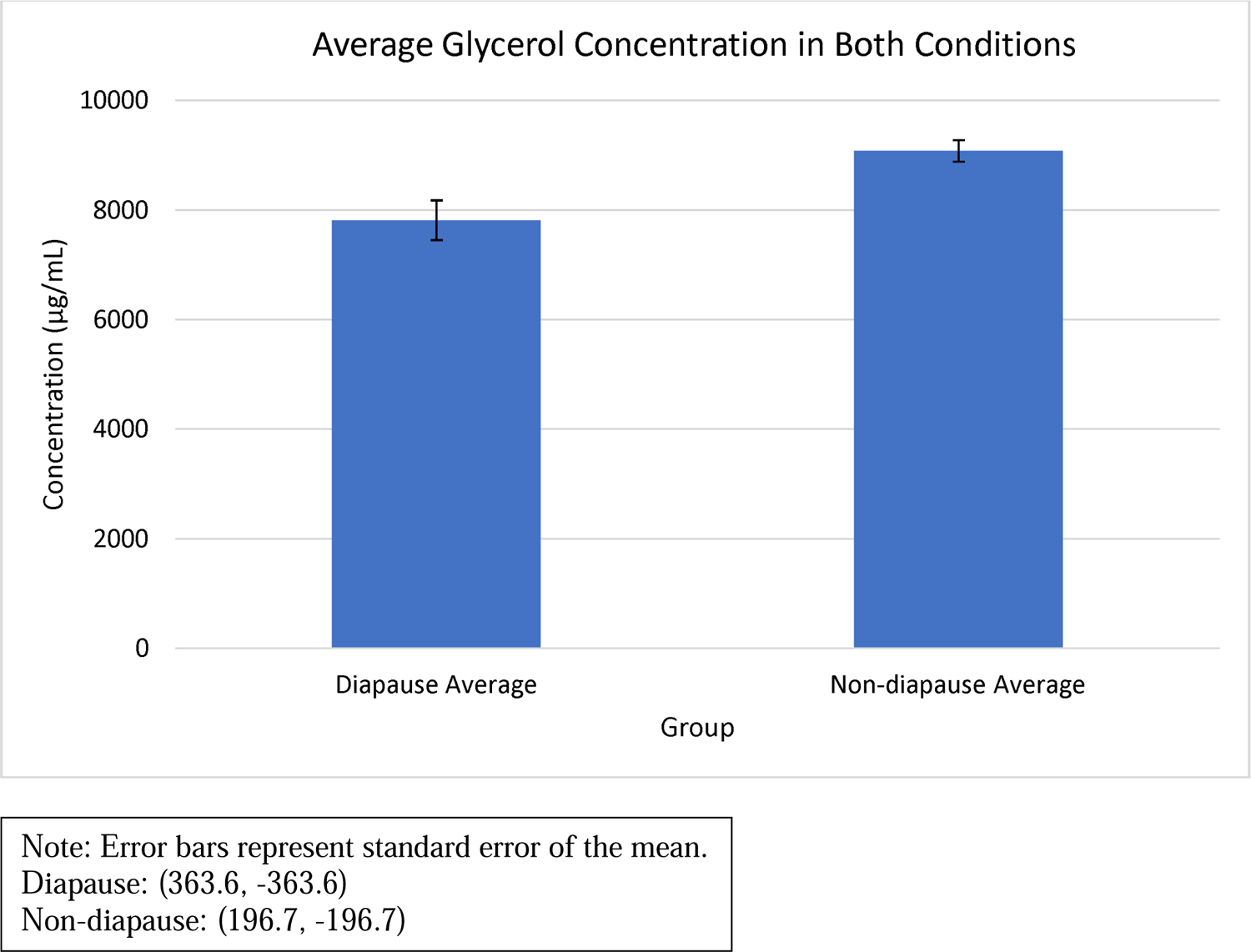
The average concentrations, in µg/mL, of the diapausing specimens and the non-diapausing specimens.

## Discussion

The results of this paper provide evidence that diapause and enhanced glycerol production are not linked in the tobacco hornworm. Interestingly, the results rejected both the null hypothesis and the alternative hypothesis, as the non-diapause specimens produced significantly higher glycerol concentrations compared to the diapause specimens. To reiterate, the alternative hypothesis claimed that the diapausing specimens will produce higher glycerol concentrations than the non-diapausing specimens. In other words, the alternative hypothesis predicted that a significant difference between the two groups would occur in the opposite direction of the obtained results. As previously stated, the most likely explanation for these results is that diapause and glycerol production are not linked in the tobacco hornworm. Most insects studied on this topic exhibit no link between diapause and glycerol upregulation. Nevertheless, this explanation does not completely explain why the non-diapausing pupae contained substantially higher glycerol concentrations compared to the diapausing pupae.

A few possible explanations may indicate why the results produced this unique significant difference. One reason is the decrease in metabolism during diapause may have decreased the glycerol concentration in the diapausing pupae. Similar to humans, glycerol is an intermediate in glycolysis (Churchill & Storey, 1989) and a byproduct of lipolysis in insects Crabtree & Newsholme, 1972). Accordingly, the decrease in metabolism associated with diapause may cause decreased glycerol concentrations since glycerol is a glycolytic intermediate and a byproduct of lipolysis. Notably, Zhai et al. (2017) observed that glycolysis was downregulated during diapause in small brown planthoppers (*Laodelphax striatellus*). Although several studies report that glycolysis is upregulated during diapause, these studies commonly expose insects to markedly cold temperatures. Therefore, the cold temperature exposure in these studies may be the underlying cause of the upregulation in glycolysis during diapause, since an upregulation in glycolysis is beneficial towards creating more glycerol for cryoprotection. For example, in one study on diapause in a parasitic wasp (*Praon volucre*), Colinet et al. (2012) denoted that glycolysis was upregulated during diapause, most probably to increase polyol synthesis. However, these researchers placed the parasitic wasps in 2°C to induce diapause (Colinet et al., 2012). Consequently, the cold temperature exposure may have been the underlying cause of the upregulation in glycolysis and increase in glycerol accumulation.

Nevertheless, photoinduction of diapause may not be possible in this species, leaving a cold temperature induced diapause as the only option. Additionally, Colinet et al.’s (2012) study is still valuable since diapause commonly occurs during cold temperature exposure in insects, even if the cold temperature exposure was not the primary cause of diapause induction.

Another possible explanation for these results is the diapause group was not given enough time to increase glycerol accumulation. In a noteworthy study on diapause in the cecropia moth (*Hyalophora cecropia*), diapausing cecropia moths accumulated greater levels of glycerol until six months of dormancy, plateauing at .3 M (Wyatt & Meyer, 1958). For reference, in the current study, diapausing tobacco hornworms contained .085 M after 16 days of diapause. Nonetheless, this temporal explanation would not explain why non-diapausing tobacco hornworms contained substantially greater concentrations of glycerol than diapausing tobacco hornworms. Accordingly, this result further supports the idea that diapause and glycerol upregulation are not linked in the tobacco hornworm. Moreover, Wyatt & Meyer (1958) initially exposed different species of diapausing insects to room temperature in the beginning months of diapause, and to cold temperatures (6–7 °C) during the later months of diapause. This was done to assess whether diapause independently influences glycerol production, or if cold temperature is also needed to upregulate glycerol production during diapause. Specifically, Wyatt & Meyer (1958) also investigated glycerol production during diapause in the tasar silkworm (*Antheraea mylitta*), Orizaba silkmoth (*Rothschildia orizaba*), polyphemus moth (*Antheraea polyphemus*), and a large Asiatic moth (*Samia walkeri*). Interestingly, out of these four insects, only the polyphemus moth displayed any meaningful glycerol presence in hemolymph during any point of diapause, at approximately .05 M. Concerning the other three related species, glycerol concentration was nearly zero in hemolymph in both the beginning months of diapause at room temperature, and the subsequent months of diapause during exposure to cold temperatures. These findings provide value to the current study, since clearly not all insects produce glycerol for cryoprotection during diapause, as these three insects displayed only trace amounts during diapause, and the polyphemus moth did not increase glycerol production with exposure to cold temperature.

Furthermore, these findings illuminate one last possible explanation for the unexpected significant difference between diapausing tobacco hornworms and non-diapausing tobacco hornworms. This final possible explanation involves the production of other cryoprotectants in response to diapause and cold temperature exposure. Although glycerol is the most commonly used cryoprotectant in insects, sorbitol, mannitol, dulcitol, ethylene glycol, threitol, inositol, sucrose, and trehalose are also used for cryoprotection in various insects (Miller & Smith, 1975; Baust & Edwards, 1979; Ring, 1982; Gehrken, 1984; Storey & Storey, 1985; Zachariassen, 1985). Notably, of all these cryoprotectants, sorbitol is the most common cryoprotectant in insects after glycerol. In a compelling experiment on this subject, Pullin & Bale (1989) revealed that sorbitol concentrations were immensely greater in diapausing cabbage butterflies (*Pieris brassicae*) compared to non-diapausing cabbage butterflies. Conversely, glycerol was only present in extremely low levels in both the diapausing and non-diapausing specimens. Pullin & Bale (1989) inferred from these findings that sorbitol is the main cryoprotectant for diapausing cabbage butterflies, while glycerol does not function as a cryoprotectant in this species. Perhaps the tobacco hornworm similarly relies more on sorbitol or other cryoprotectants instead of glycerol during diapause. Therefore, upregulation of sorbitol or other cryoprotectants may be linked to diapause in the tobacco hornworm, as the insect may rely more on cryoprotectants other than glycerol. Ultimately, although the tobacco hornworm may depend on cryoprotectants other than glycerol, a link between the production of these cryoprotectants and diapause may still not exist.

Clearly, more research needs to be performed on this topic to completely understand diapause and cryoprotection, and whether any link exists between the two elements in insects. Regarding the tobacco hornworm, a similar study to the current study should be conducted to examine whether more time is needed for the diapausing pupae to accumulate glycerol. This experiment would rule out the possible explanation that not enough time was provided for glycerol accumulation in the diapausing specimens. Another experiment that should be conducted in the tobacco hornworm would use the same methodology as the current study, except with four groups instead of two groups. The additional two groups would be diapausing pupae and non-diapausing pupae that are exposed to cold temperature during rearing. The results of this study would uncover whether cold temperatures can exclusively elicit glycerol upregulation, or if diapause and cold temperature exposure are needed in combination to cause glycerol upregulation. Of course, one more possible result is that neither cold temperature nor diapause influences glycerol production in the tobacco hornworm. This would probably be the case if the tobacco hornworm does not rely on glycerol for cryoprotection, but is dependent on other biomolecules instead. Lastly, more experiments should be performed on this topic concerning sorbitol and other cryoprotectants to analyze whether a linkage between diapause and these other cryoprotectants is evident.

## Acknowledgements

Words cannot express my gratitude to my professor and supervisor, Dr. Julian Shepherd, for his teachings and invaluable feedback for my research. My journey into entomology would have never happened without him and his incredible teachings. His class was an inspiration to my research on this topic and my continued passion for research upon graduating from Binghamton University.

## References

Araújo, P. M., Viegas, I., Rocha, A. D., Villegas, A., Jones, J. G., Mendonça, L., Ramos, J. A., Masero, J. A., & Alves, J. A. (2019). Understanding how birds rebuild fat stores during migration: insights from an experimental study. Scientific Reports, 9(1), 1–11. 10.1038/s41598-019-46487-z

Baust, J. G., & Edwards, J. S. (1979). Mechanisms of freezing tolerance in an Antarctic midge, Belgica antarctica. Physiological Entomology, 4(1), 1–5. 10.1111/j.1365-3032.1979.tb00171.x

Bell, R. A., & Joachim, F. G. (1976). Techniques for Rearing Laboratory Colonies of Tobacco Hornworms and Pink Bollworms. Annals of the Entomological Society of America, 69(2), 365–373. 10.1093/aesa/69.2.365

Bell, R. A., Rasul, C. G., & Joachim, F. G. (1975). Photoperiodic induction of the pupal diapause in the tobacco hornworm, Manduca sexta. Journal of Insect Physiology, 21(8), 1471–1480. 10.1016/0022-1910(75)90210-3

Berkley, C. (2019). Teach Life Cycles with the Tobacco Hornworm | Carolina.com.

Carolina.Com. https://www.carolina.com/teacher-resources/Interactive/teach-life-cycles-with-the-tobacco-hornworm/tr30179.tr

Byron, M., & Gillet-Kaufman, J. (2017). tomato hornworm, Manduca quinquemaculata (Haworth). Ufl.Edu. http://entnemdept.ufl.edu/creatures/field/hornworm.htm

Chanda, B., Venugopal, S. C., Kulshrestha, S., Navarre, D. A., Downie, B., Vaillancourt, L., Kachroo, A., & Kachroo, P. (2008). Glycerol-3-phosphate levels are associated with basal resistance to the hemibiotrophic fungus Colletotrichum higginsianum in Arabidopsis. Plant Physiology, 147(4), 2017–2029. 10.1104/pp.108.121335

Churchill, T. A., & Storey, K. B. (1989). Metabolic correlates to glycerol biosynthesis in a freeze-avoiding insect,Epiblema scudderiana. Journal of Comparative Physiology B, 159(4), 461–472. 10.1007/bf00692418

Colinet, H., Renault, D., Charoy-Guével, B., & Com, E. (2012). Metabolic and Proteomic Profiling of Diapause in the Aphid Parasitoid Praon volucre. PLoS ONE, 7(2), e32606. 10.1371/journal.pone.0032606

Crabtree, B., & Newsholme, E. A. (1972). The activities of lipases and carnitine palmitoyl-transferase in muscles from vertebrates and invertebrates. Biochemical Journal, 130(3), 697–705. https://www.ncbi.nlm.nih.gov/pmc/articles/PMC1174508/

Denlinger, D. (2009). Diapause - an overview | ScienceDirect Topics. www.Sciencedirect.Com. https://www.sciencedirect.com/topics/agricultural-and-biological-sciences/diapause

Dubach, P., Pratt, D., Smith, F., & Stewart, C. M. (1959). Possible Role of Glycerol in the Winter-Hardiness of Insects. Nature, 184(4682), 288–289. 10.1038/184288b0

Frankos, V. H., & Platt, A. P. (1976). Glycerol accumulation and water content in larvae of Limenitis archippus: Their importance to winter survival. Journal of Insect Physiology, 22(5), 623–628. 10.1016/0022-1910(76)90225-0

Fraser, J. D., Bonnett, T. R., Keeling, C. I., & Huber, D. P. W. (2017). Seasonal shifts in accumulation of glycerol biosynthetic gene transcripts in mountain pine beetle, Dendroctonus ponderosae Hopkins (Coleoptera: Curculionidae), larvae. PeerJ, 5(e3284). 10.7717/peerj.3284

Gehrken, U. (1984). Winter survival of an adult bark beetle Ips acuminatus Gyll. Journal of Insect Physiology, 30(5), 421–429. 10.1016/0022-1910(84)90100-8

Gerber, D. W., Byerrum, R. U., Gee, R. W., & Tolbert, N. E. (1988). Glycerol concentrations in crop plants. Plant Science, 56(1), 31–38. 10.1016/0168-9452(88)90182-3

Grasso, R., Musumeci, F., Gulino, M., & Scordino, A. (2018). Exploring the behaviour of water in glycerol solutions by using delayed luminescence. PLoS ONE, 13(1). 10.1371/journal.pone.0191861

Hanec, W., & Beck, S. D. (1960). Cold hardiness in the European corn borer, Pyrausta nubilalis (Hübn.). Journal of Insect Physiology, 5(3), 169–180. 10.1016/0022-1910(60)90002-0

Ishiguro, S., Li, Y., Nakano, K., Tsumuki, H., & Goto, M. (2007). Seasonal changes in glycerol content and cold hardiness in two ecotypes of the rice stem borer, Chilo suppressalis, exposed to the environment in the Shonai district, Japan. Journal of Insect Physiology, 53(4), 392–397. 10.1016/j.jinsphys.2006.12.014

Lee, R. E., Chen, C., Meacham, M. H., & Denlinger, D. L. (1987). Ontogenetic patterns of cold-hardiness and glycerol production in Sarcophaga crassipalpis. Journal of Insect Physiology, 33(8), 587–592. 10.1016/0022-1910(87)90074-6

Leithner, K., Triebl, A., Trötzmüller, M., Hinteregger, B., Leko, P., Wieser, B. I., Grasmann, G., Bertsch, A. L., Züllig, T., Stacher, E., Valli, A., Prassl, R., Olschewski, A., Harris, A. L., Köfeler, H. C., Olschewski, H., & Hrzenjak, A. (2018). The glycerol backbone of phospholipids derives from noncarbohydrate precursors in starved lung cancer cells. Proceedings of the National Academy of Sciences of the United States of America, 115(24), 6225–6230. 10.1073/pnas.1719871115

Miller, L. K., & Smith, J. S. (1975). Production of threitol and sorbitol by an adult insect: association with freezing tolerance. Nature, 258(5535), 519–520. 10.1038/258519a0

Nelson, J. L., Harmon, M. E., & Robergs, R. A. (2011). Identifying plasma glycerol concentration associated with urinary glycerol excretion in trained humans. Journal of Analytical Toxicology, 35(9), 617–623. 10.1093/anatox/35.9.617

Pullin, A. S., & Bale, J. S. (1989). Influence of diapause and temperature on cryoprotectant synthesis and cold hardiness in pupae of Pieris brassicae. Comparative Biochemistry and Physiology Part A: Physiology, 94(3), 499–503. 10.1016/0300-9629(89)90128-X

Rickards, J., Kelleher, M. J., & Storey, K. B. (1987). Strategies of freeze avoidance in larvae of the goldenrod gall moth, Epiblema scudderiana: Winter profiles of a natural population. Journal of Insect Physiology, 33(6), 443–450. 10.1016/0022-1910(87)90024-2

Ring, R. A. (1982). Freezing-tolerant insects with low supercooling points. Comparative Biochemistry and Physiology Part A: Physiology, 73(4), 605–612. 10.1016/0300-9629(82)90267-5

Rotondo, F., Ho-Palma, A. C., Remesar, X., Fernández-López, J. A., Romero, M. del M., & Alemany, M. (2017). Glycerol is synthesized and secreted by adipocytes to dispose of excess glucose, via glycerogenesis and increased acyl-glycerol turnover. Scientific Reports, 7(8983). 10.1038/s41598-017-09450-4

Storey, J. M., & Storey, K. B. (1986). Winter survival of the gall fly larva, Eurosta solidaginis: Profiles of fuel reserves and cryoprotectants in a natural population. Journal of Insect Physiology, 32(6), 549–556. 10.1016/0022-1910(86)90070-3

Sturgeon, R. J., Deamer, R. L., & Harbison, H. A. (1979). Improved Spectrophotometric Determination of Glycerol and Its Comparison with an Enzymatic Method. Journal of Pharmaceutical Sciences, 68(8), 1064–1066. 10.1002/jps.2600680841

Wood, F. E., & Nordin, J. H. (1976). Studies on the low temperature induced biogenesis of glycerol by adult Protophormia terranovae. Journal of Insect Physiology, 22(12), 1665–1673. 10.1016/0022-1910(76)90060-3

Woude, H. A. v. d., & Verhoef, H. A. (1988). Reproductive diapause and cold hardiness in temperate Collembola Orchesella cincta and Tomocerus minor. Journal of Insect Physiology, 34(5), 387–392. 10.1016/0022-1910(88)90108-4

Wyatt, G. R., & Meyer, W. L. (1959). THE CHEMISTRY OF INSECT HEMOLYMPH. The Journal of General Physiology, 42(5), 1005–1011. https://www.ncbi.nlm.nih.gov/pmc/articles/PMC2194945/

Yang, Y., Zhao, J., Liu, P., Xing, H., Li, C., Wei, G., & Kang, Z. (2013). Glycerol-3-Phosphate Metabolism in Wheat Contributes to Systemic Acquired Resistance against Puccinia striiformis f. sp. tritici. PLoS ONE, 8(11), e81756. 10.1371/journal.pone.0081756

Zachariassen, K. E. (1985). Physiology of cold tolerance in insects. Physiological Reviews, 65(4), 799–832. 10.1152/physrev.1985.65.4.799

Zhai, Y., Zhang, Z., Gao, H., Chen, H., Sun, M., Zhang, W., Yu, Y., & Zheng, L. (2017). Hormone Signaling Regulates Nymphal Diapause in Laodelphax striatellus (Hemiptera: Delphacidae). Scientific Reports, 7(1), 1–9. 10.1038/s41598-017-13879-y

